# Manganese-enhanced MRI (MEMRI) in breast and prostate cancers: exploring the potential role of calcium receptors

**DOI:** 10.1101/806448

**Authors:** Gabriella Baio, Marina Fabbi, Michele Cilli, Francesca Rosa, Simona Boccardo, Francesca Valdora, Sandra Salvi, Luca Basso, Laura Emionite, Eliana Gianolio, Silvio Aime, Carlo Emanuele Neumaier

## Abstract

**Procedures:** To assess the relationship between Manganese Enhanced Magnetic Resonance Imaging (MEMRI) and the expression of calcium receptors in human prostate and breast cancer animal models.

**Methods:** NOD/SCID mice were inoculated with MDA-MB-231 breast cancer cells and prostate PC3 cancer cells to develop orthotopic or pseudometastatic cancer animal models. Mice were studied on a clinical 3T scanner by using a prototype birdcage coil before and after intravenous injection of MnCl_2_. Assessment of receptor’s status was carried out after the MR images acquisition by immunohistochemistry on excised tumours.

**Results:** Manganese contrast enhancement in breast or prostate cancer animal models well correlated with CaSR expression (p<0.01), whereas TRPV6 expression levels appeared not relevant to the Mn uptake.

**Conclusion:** MEMRI appears an efficient tool to characterize human breast and prostate cancer animal models in the presence of different expression level of CaSR.

## Introduction

Regulation of calcium metabolism is crucial for cellular activities such as proliferation, gene transcription and cell death (1, 2). To maintain a balanced condition between the intra- and extra-cellular compartments, cells employ specialized calcium pumps, channels and calcium binding proteins (the so called “molecular toolkit”) (3, 4). Changes in calcium metabolism are reported on the upsurge of certain pathological states, including cancer (5–7). To better detect the occurrence of an altered calcium metabolism *in vivo*, a non-invasive method such as imaging could be extremely valuable. On the basis of the close analogy between calcium and manganese, an imaging technique such as Manganese Enhanced Magnetic Resonance Imaging (MEMRI) has received increased attention in preclinical cancer imaging studies (8–10).

MEMRI is a well-established method that, by combining the peculiar physical and biological properties of Mn^2+^, allows to extract relevant anatomical and functional imaging information in a variety of systems (11–14).

Previously we tested MEMRI as imaging tool to report on the high expression levels of Calcium Sensing Receptor (CaSR) in a human breast cancer animal model (15). CaSR is a G protein coupled receptor, which plays an important role in tumour development (16–19). Breast tumours displaying high CaSR expression, as assessed by immunohistochemistry, correlated well with manganese uptake assessed by Magnetic Resonance Imaging (MRI). However, CaSR is not the only receptor involved in the presence of the altered calcium metabolism in cancer cells, as other calcium receptors or channels can interfere with Ca^2+^ homeostasis. Calcium-permeable ion channels, such as Transient Receptor Potential (TRP) have been recently associated with tumour progression (20). An altered expression level of TRP channels is associated with changes in intracellular Ca^2+^ and in the proliferative pathways, promoting or inhibiting apoptosis of the tumour cells (21, 22). Specifically, TRPV6 is a highly Ca^2+^-selective channel (23–25) and its activity is also known to be modulated by oestrogen, progesterone, tamoxifen and vitamin D, affecting proliferation and survival of cancer cells (26). In prostate and breast cancer, TRPV6 is substantially associated with highly proliferative and metastatic tumours (26, 27). A high level of expression of TRPV6 is associated with high Gleason score tumours and high metastatic risk (28, 29), supporting its potential use to predict the clinical outcome of human prostate cancer (30). Similarly, in oestrogen receptor-negative breast cancer, high TRPV6 levels were associated to decreased survival rates when compared to patients with low or intermediate TRPV6 expression (31).

Considering the important role that TRPV6 and CaSR have in cancer, we undertook these pilot experiments to investigate the MEMRI response in different types of breast and prostate cancer animal models, characterized by different level of expression of these molecules.

## Results

To elucidate the relationship between the cellular uptake of Mn^2+^ and CaSR and TRPV6 expression, different murine models have been prepared, namely human breast orthotopic model, human prostate orthotopic model, intraosseous human breast model and human pseudometastatic prostate model. For each model, after the acquisition of the MEMRI experiment, the animals were sacrificed and the tumour cells analyzed for assessing the level of expression of CaSR and TRPV6, respectively.

### Human breast orthotopic cancer animal model

MDA-MB-231 breast cancer cells were inoculated into NOD-SCID mice (n=4) and MR images were acquired starting from week 6 after the cells transplant. T_2W_-MR imaging showed a rounded, well defined, lesion within the mammary fat-pad without any sign of infiltration of the surrounding tissue. The dynamics of manganese uptake resulted quite different among the considered mice. In two mice, 10 minutes after manganese administration, a slight peripheral ring of contrast enhancement was observed in the tumour region, while a homogeneous contrast enhancement was gradually appreciated at later time points (30, 60 and 90 minutes) (Figure 1 AB). Quantitative analysis demonstrated an increased tumour signal enhancement (SE) of 91% at 90 min from the Mn^2+^ administration (see Table 1). In the other two mice, no manganese uptake was observed at all-time points (Figure 1 EF) as witnessed by the quantitative analysis that demonstrated no significant changes in SE (see Table 1).

**Figure 1.**
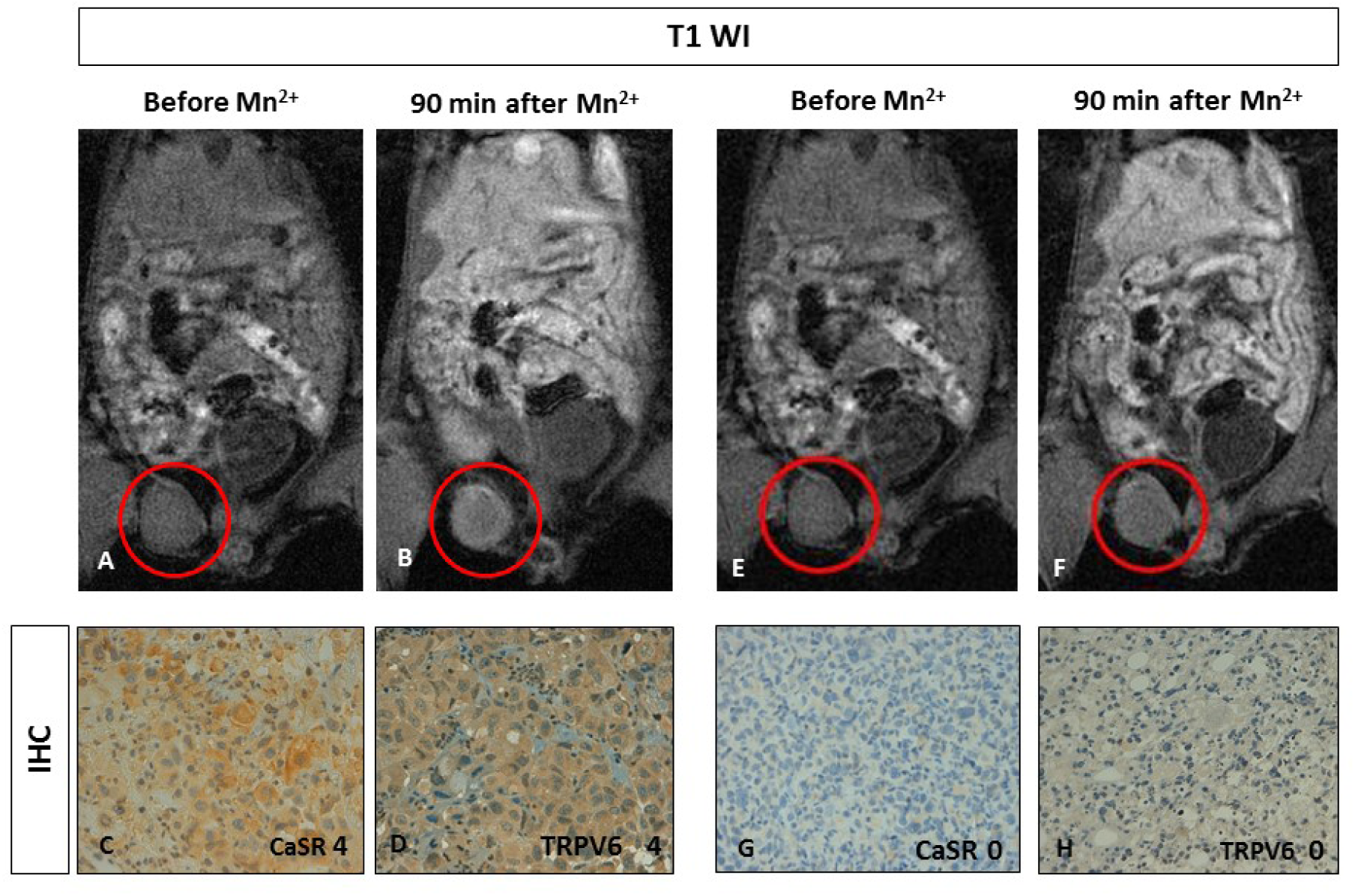
Manganese enhanced MR imaging (MEMRI) of orthotopic human breast cancer xenografts and CaSR/TRPV6 receptors levels. **AB.** Tumour diameter 5 mm. Upon comparing T1-weighted gradient echo images (T1 WI) recorded before and 90 minutes after Mn^2+^ administration, the tumour showed manganese uptake (red circles). **CD.** Immunohistochemistry (IHC) of CaSR and TRPV6: both receptors are detected (score 4) with a homogenous intense staining. Original magnification 200x. **EF.** Tumour diameter 5 mm. Upon comparing T1-weighted gradient echo images (T1 WI) recorded before and 90 minutes after Mn2+ administration, no changes in tumour signal enhancement was appreciated (red circles). **GH.** Immunohistochemistry (IHC) showed no evidence for CaSR and TRPV6 expression (score 0). Original magnification 200x.

**Table 1.** SNR and SE% (calculated as [(SNR-SNR^0^)/ SNR^0^]×100) for all tumour lesions before and 90 minutes after manganese administration and CaSR or TRPV6 expression levels evaluated by immunohistochemistry.

|  | Before | After Mn<sup>2+</sup> | SE | CaSR | TRPV6 |
| --- | --- | --- | --- | --- | --- |
| **Xenotransplant lesions** | **SNR** | **SNR** | **%** | **score** | **Score** |
| Orthotopic PC3 | 7,61 | 7,64 | +0.4 | 0 | 0 |
| Orthotopic PC3 | 4,90 | 5,27 | +7.5 | 1 | 1 |
| Orthotopic MDA-MB-231 | 5,99 | 5,87 | -2.0 | 0 | 0 |
| Orthotopic MDA-MB-231 | 6,15 | 11,75 | +91 | 4 | 4 |
| Osseous MDA-MB-231 | 10,11 | 9,35 | -7.5 | 0 | 0 |
|  | Before | After Mn<sup>2+</sup> | SE | CaSR | TRPV6 |
| **Pseudo-metastatic lesions (PC3)** | **SNR<sup>0</sup>** | **SNR** | **%** | **score** | **score** |
| Liver nodule | 11,28 | 10,98 | -2.6 | 1 | 4 |
| Liver nodule | 14,52 | 13,70 | -5.6 | 1 | 4 |
| Mediastinal nodule | 11,13 | 14,85 | +33 | 3 | 2 |
| Mediastinal nodule | 11,48 | 13,41 | +17 | 2 | 2 |
| Diaphragmatic nodule | 8,97 | 13,15 | +47 | 3 | 4 |
| Peritoneal nodule | 11,82 | 11,77 | -0.4 | 1 | 1 |
| Peritoneal nodule | 13,76 | 13,64 | -0.9 | 1 | 1 |

**Table 2.**
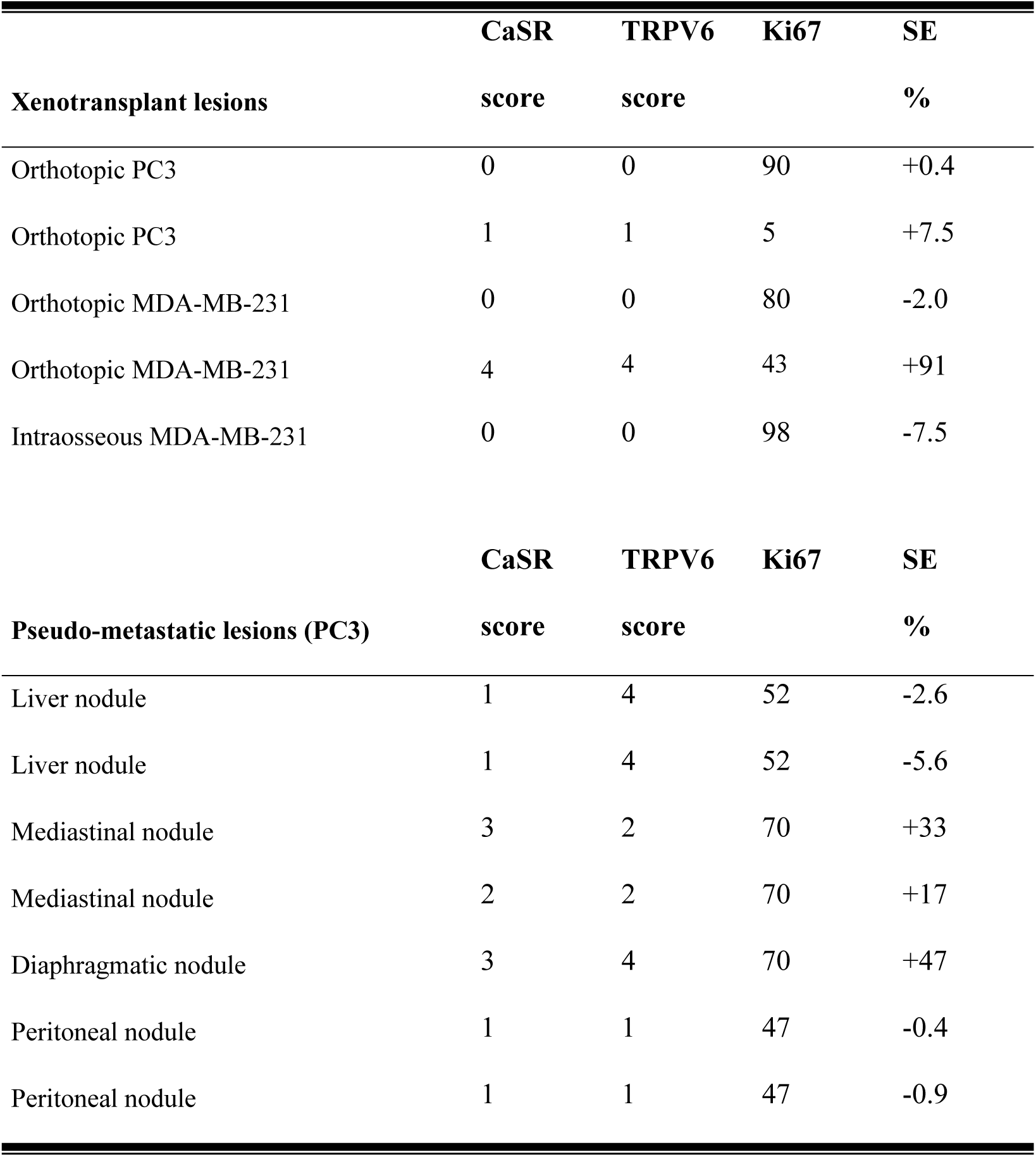
CaSR, TRPV6 and Ki67 status of each lesion between the different type of human breast or prostate cancer animal models.

To correlate manganese-enhancement with CaSR or TRPV6 expression, immunohistochemistry of the excised lesions was performed. As shown in Figure 1 CD and GH, in these tumours the expression levels of CaSR and TRPV6 ranged between score 0 and 4 and both calcium receptors appear expressed at the same level in the same tumour. Interestingly, the CaSR or TRPV6 score levels correlated well with MEMRI SI observations.

### Human prostate orthotopic cancer animal model

The orthotopic prostate xenotransplant model (n=2) was prepared by implantation of PC3 cells in the prostate gland tissue under sterile surgical conditions. T_2W_-MR images showed well-defined lobulated lesions in both animals after 6 weeks (Figure 2 AF). No manganese enhancement was detected within both tumours at all acquisition time points (10, 30 and 90 minutes) (Figure 2 BC, GH). The quantitative analysis showed no significant difference in SE before and after manganese administration in both animals (see table 1). The immunohistochemistry analysis of prostate tumours demonstrated a very low expression level of CaSR and TRPV6 with a score between 0 and 1 (Figure 2 DE, IL). The level of expression of both receptors was consistent in the same tumour.

**Figure 2.**
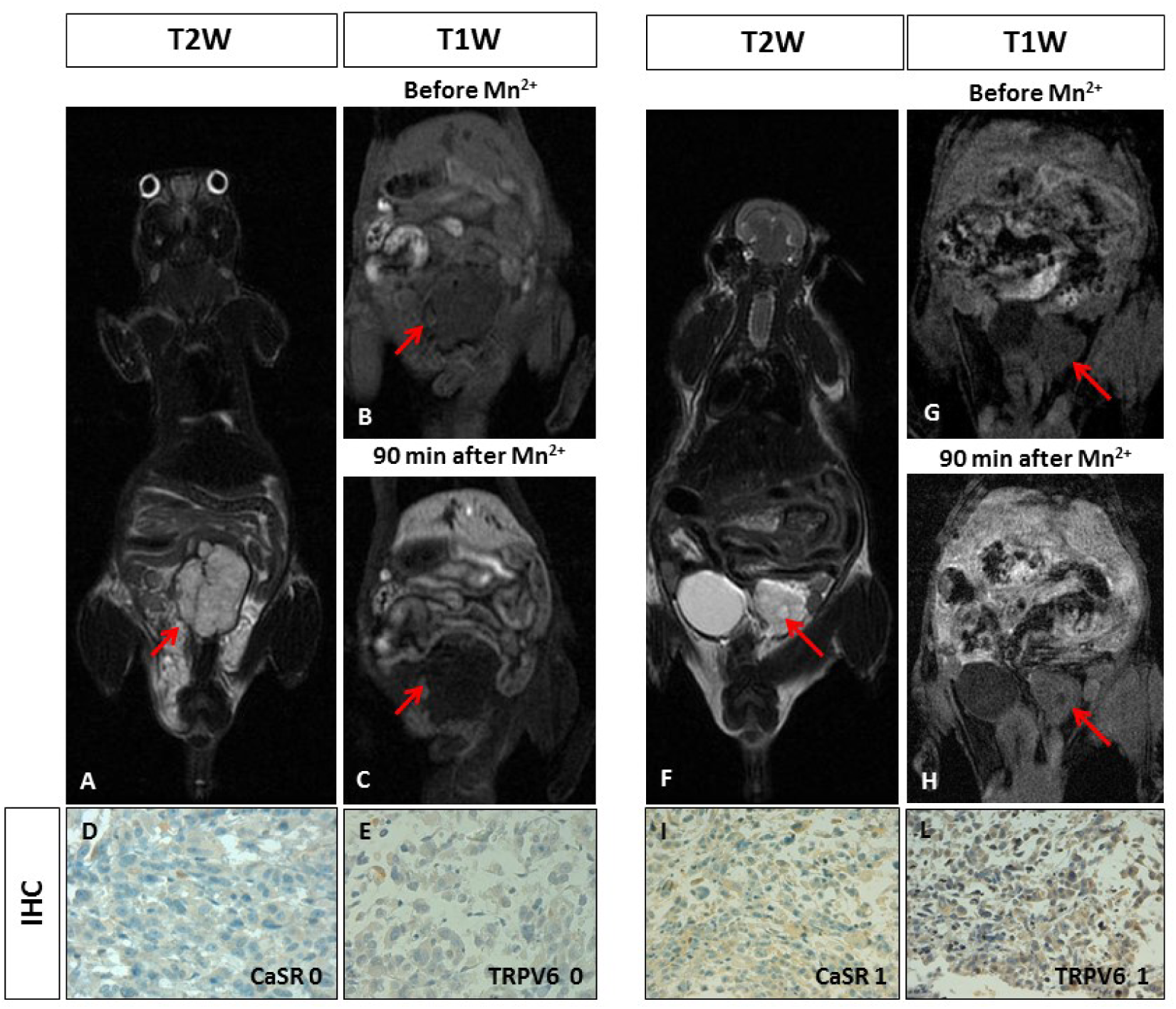
Manganese enhanced MR imaging (MEMRI) of orthotopic human prostate cancer xenografts and CaSR/TRPV6 receptors levels. Tumour diameter 10 mm. On T2-weighted MR image a high signal intensity multilobulated prostatic tumour is detected (red arrow). **BC.** On T1-weighted gradient echo MR images (T1 WI) recorded before and 90 minutes after Mn^2+^ administration, no changes in tumour signal enhancement was appreciated (red arrows). **DE.** Immunohistochemistry (IHC) of CaSR and TRPV6: both receptors were not detected in tumour cells (score 0) showing a negative staining. Original magnification 200x. **F.** Tumour diameter 6 mm. On T2-weighted image a high signal intensity extra-capsular prostatic tumour is detected (red arrow). **GH.** On T1-weighted gradient echo images (T1 WI) recorded before and 90 minutes after Mn2+ administration, no changes in tumour signal enhancement was appreciated (red arrows). **IL.** Immunohistochemistry (IHC) of CaSR and TRPV6: both receptors displayed a very low positive staining in tumour cells (score 1). Original magnification 200x.

### Intraosseous human breast cancer xenotransplant model

To investigate the manganese uptake in pseudo-metastatic bone breast cancer lesions, the intraosseous human breast cancer xenotransplant model was used. The osseous cancer model was set up according to Corey et al. (32). As shown in figure 3, T2 weighted MR images of the bone xenotransplant appeared as a well-defined soft rounded lesion in the left-leg (Figure 3AE). Radiographic examination showed a cortical swelling of the tibia and tiny lytic lesions (Figure 3D). Upon manganese administration, on T1-weighted images, no contrast enhancement was observed within the tumour mass at all acquisition time points (Figure 3BC) as confirmed by the quantitative analysis (see table 1). The immunohistochemistry analysis showed no CaSR and TRPV6 (Figure 3 FG) expression.

**Figure 3.**
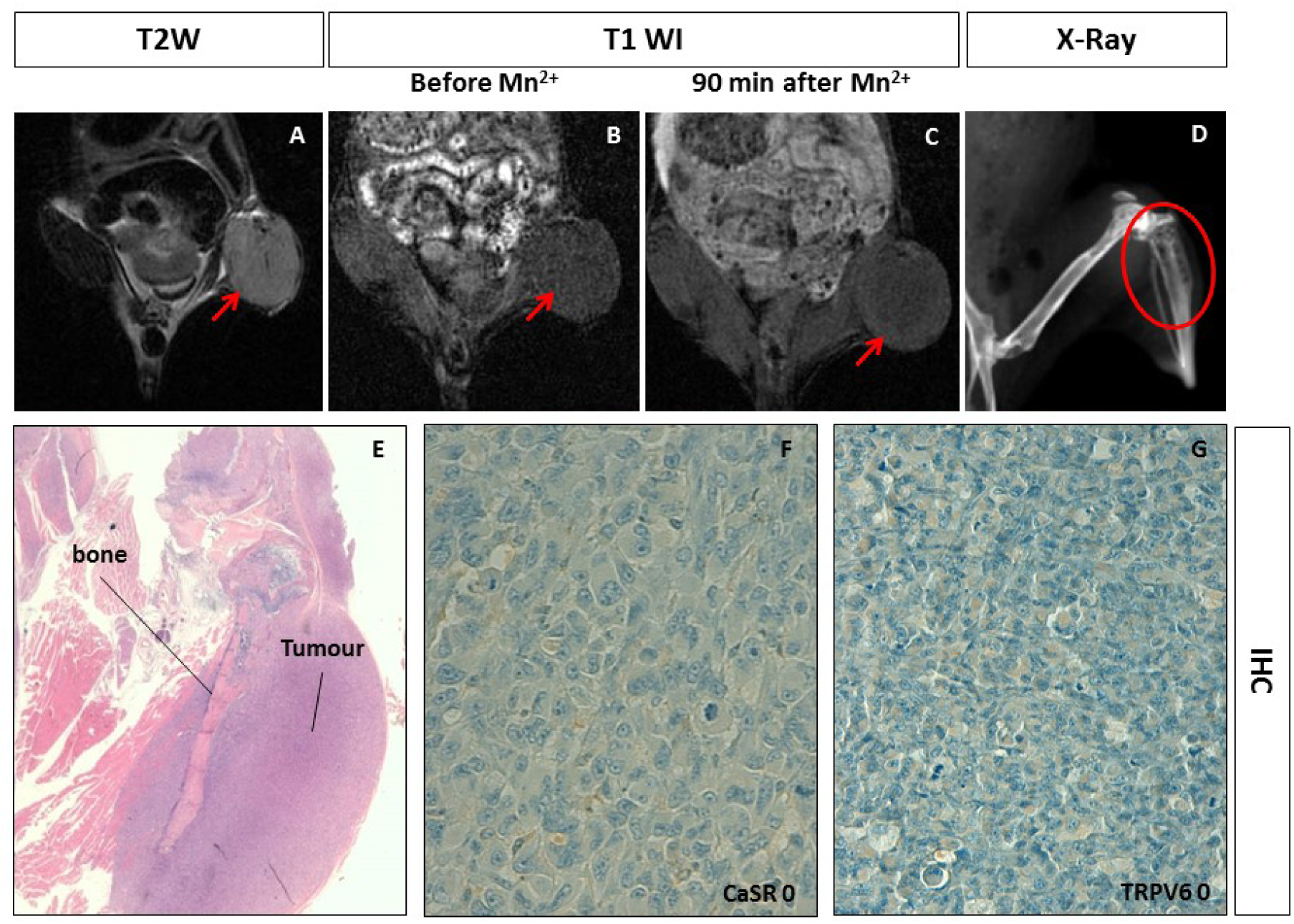
Manganese enhanced MR imaging (MEMRI) of the intraosseous human breast cancer xenotransplant model and CaSR/TRPV6 receptors levels. **A.** Tumor diameter 10 mm. On T2-weighted MR image a low signal intensity solid tumor is detected at the left leg of the mouse (red arrow). **BC.** On T1-weighted gradient echo MR images (T1 WI) recorded before and 90 minutes after Mn^2+^ administration, no changes in tumor enhancement was appreciated (red arrows). **D.** X-Ray imaging of the left leg of the mouse showed multiple lytic areas within the tibia in keeping with the tumor transplantation (red circle). **E.** Hematoxylin and eosin staining displaying the intraosseous tumor. **FG.** Immunohistochemistry (IHC) of CaSR and TRPV6: both receptors were not detected in tumor cells (score 0) showing a negative staining. Original magnification 200x.

### Human pseudometastatic prostate cancer animal model

The pseudometastatic prostate cancer animal models, obtained by intra-cardiac injection of PC3 cells, developed multiple lesions in the mediastinum, liver and peritoneal deposits with ascites, respectively (Figure 4A and 5A). After 90 min, the prostatic lesions showed different manganese uptake (from peripheral to full contrast enhancement) which remained stable for 3 hours (Figure 4 B-Q and 5B-C). Quantitative analysis demonstrated a large heterogeneity in the SE of all the metastatic deposits (SI range from −0.9 to 47% (see table 1). Different expression levels of both calcium receptors (ranging from 0 to 3 for CaSR and from 0 to 4 for TRPV6) was observed, in particular, in some of the prostatic lesions the expression levels of the two calcium receptors were different (Figure 4 D-S and 5D-G).

**Figure 4.**
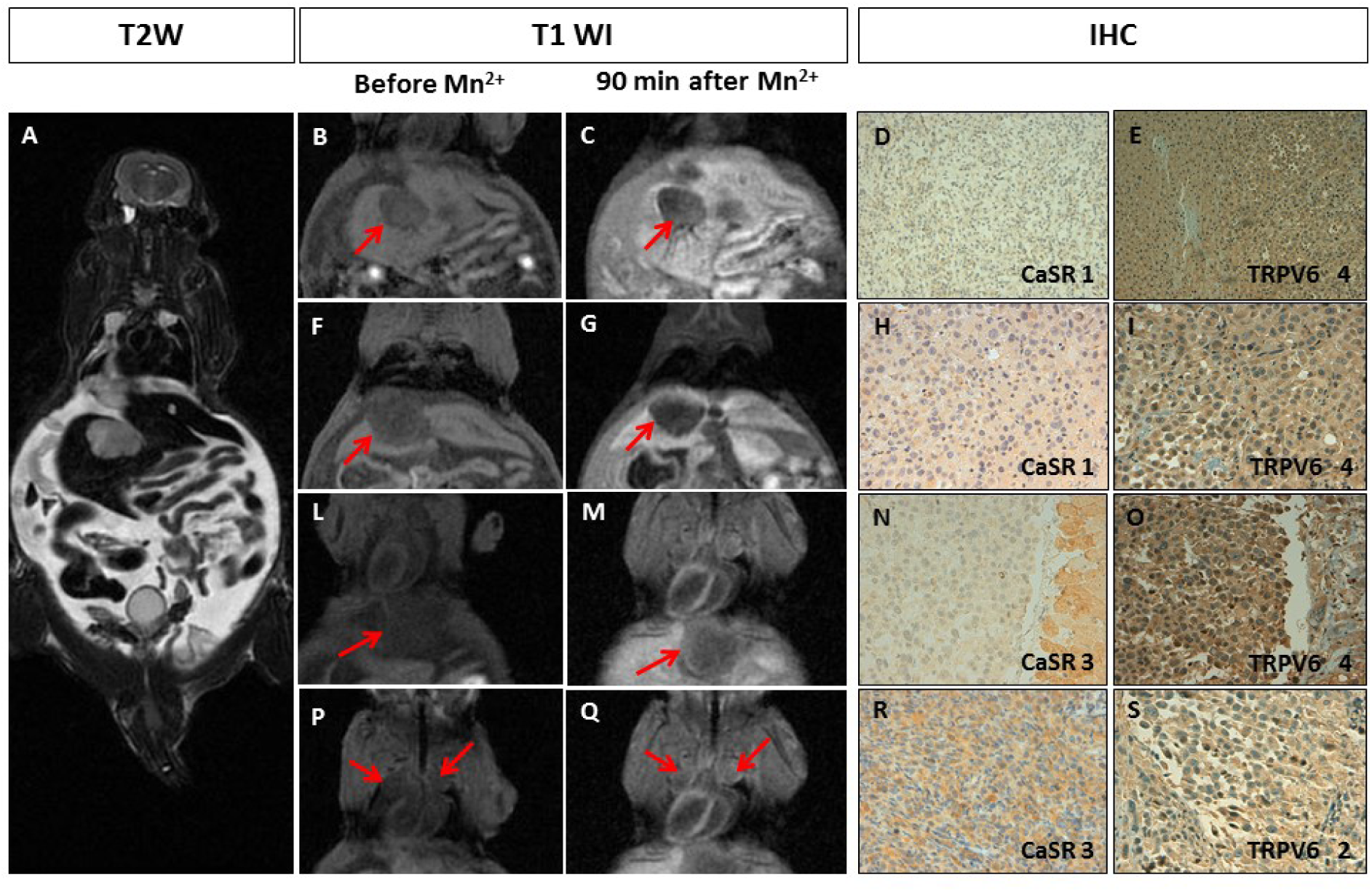
Manganese enhanced MR imaging (MEMRI) of the pseudometastatic prostate cancer animal model and CaSR/TRPV6 receptors levels. **A.** Multiple tumours with diameter from 3 to 10 mm. T2-weighted MR images showed multiple high intensity nodules with intra-abdominal ascites. **B-Q.** T1-weighted gradient echo MR images (T1 WI) recorded before and 90 minutes after Mn^2+^ administration: **B-G.** No manganese uptake by the liver metastases was detected (red arrows); **L-Q.** An increase in signal enhancement after manganese administration, respectively, within a diaphragmatic (LM) and mediastinal nodules (PQ) is appreciated (red arrows). **D-S.** Immunohistochemistry (IHC) of CaSR and TRPV6: **D-I.** A rare positive staining of CaSR was detected in tumour cells (score 1), while TRPV6 receptors displayed intense staining (score 4); **N-S.** Non-uniform weak/intense expression of CaSR was detected in tumour cells (score 3), while TRPV6 receptors ranged from strong uniform (score 4) to rare positive (score 2). Original magnification 200x.

**Figure 5.**
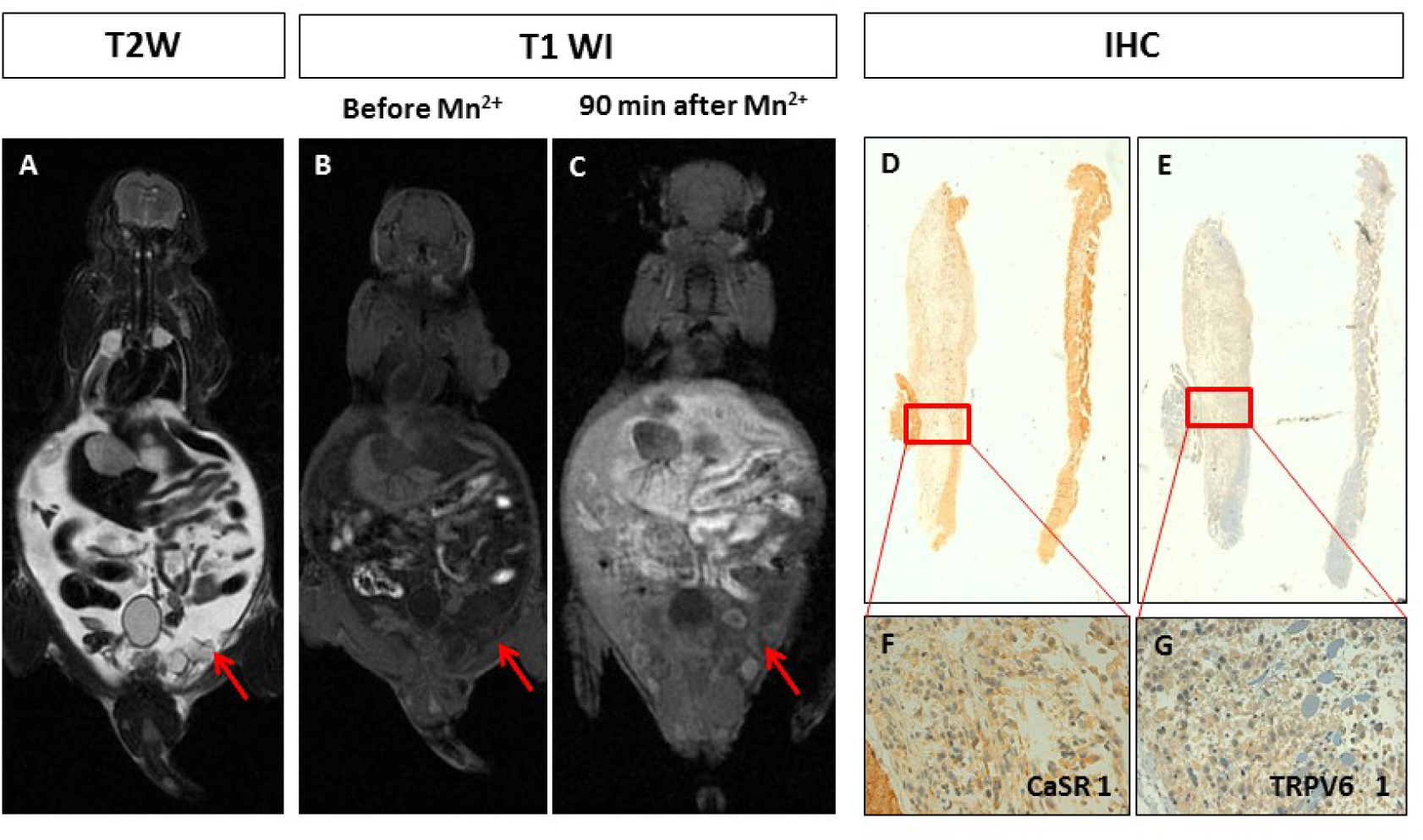
Manganese enhanced MR imaging (MEMRI) of pseudo-metastatic prostate cancer animal model and CaSR/TRPV6 receptors levels. **A.** Multiple peritoneal nodules with diameter of 3 mm (the same mouse of figure 4). In T2-weighted MR image multiple high intensity peritoneal nodules with ascites are detected. **BC.** This slice shows the peritoneal nodules in T1-weighted gradient echo images (T1 WI) recorded before and 90 minutes after Mn^2+^ administration. No significant manganese uptake is detected (red arrows); **D-G.** Immunohistochemistry (IHC) of CaSR and TRPV6: A rare positive staining of both receptors was detected in tumour cells (score 1). Original magnification 200x.

## Discussion

Several studies demonstrated the important role of calcium channels in carcinogenesis and in the modulation of cancer progression by inducing high metastatic risk (33–36). Thus, it was deemed important to explore the use of non-invasive imaging methods, to better investigate these receptors and their relationship with cancer development.

In our previous study, we applied MEMRI to image breast cancer tumours with a high-level expression of CaSR (15). Calcium sensing receptor is associated with cancer cell proliferation and bone metastatic risk in breast cancer patients (16, 17). In the present work, we expanded the study to Transient Receptor Potential channels (TRP) that are widely expressed both in normal tissue and cancer (20). Among all TRP-channels, TRPV6 is a highly Ca^2+^-selective channel and therefore quite distinguishable especially in Ca^2+^-related intracellular pathways.

We used MDA-MB-231 breast and PC3 prostate cancer cell lines normally expressing both CaSR and TRPV6 at high levels (data not shown). However, in *in vivo* cancer animal models, these tumour cells displayed different expression levels (score 0-4) for both calcium receptors. Different manganese uptake was observed in the corresponding MR images. Orthotopic human breast tumours showed either absence (score from 0) or high-level expression (score 4) of both receptors at the immunohistochemistry analysis, while in the orthotopic human prostate tumours both receptors were low expressed or absent (score 0-1). Manganese uptake appeared well correlated with the level of expression of both receptors (Figure 1 and 2).

The different expression level (score 0-4) of calcium receptors observed in *in vivo* was not expected, since these cancer cell lines are expressing high level of both receptors. The mechanism leading to the “presence or absence” of these receptors in tumour is unclear. In our previous clinical studies, we demonstrated a different CaSR level expression (score 0-4) in human primary breast cancers (36), in keeping with other clinical studies that further correlated the different expression level of CaSRs to the tumour proliferation and, in turn, to high risk of bone metastases (35).

The orthotopic breast or prostate cancer animal model results are suggesting, at a first glance, a potential combined role of both calcium receptors in tumour manganese uptake. Previous in vitro studies demonstrated the link between CaSR and TRPV6, where TRPV6 action induces calcium ions entry in CaSR (37, 38). The potential leading role of calcium receptors in manganese uptake has also been recently demonstrated by Castets CR et al, in a study where breast or glioma cancer cells lines pre-treated with Mn^2+^ and then incubated with calcium blockers enabled to retain Mn^2+^ ions into the cells. (39).

Metastatic cancer animal models are very challenging due to the reproducibility of invasiveness of the models; moreover, they showed to recapitulate the intra and inter-tumour heterogeneity usually observed in human metastatic tumours (bone and visceral metastases). We developed intraosseous human breast cancer animal models (for bone metastasis) and metastatic prostate cancer animal models (for visceral metastases) and investigated the role of MEMRI in these types of cancer animal models. The intraosseous human breast cancer animal models did not show manganese uptake in keeping with the quantitative analysis of SE. SE changes at MEMRI well correlated with both calcium receptors status detected by immunohistochemistry (CaSR and TRPV6 score 0). In these models, MEMRI and receptors expression levels behaved analogously to what observed in the orthotopic breast or prostate cancer animal models.

It is worth of notice that the two mice with metastatic prostate cancer developed visceral metastases at very different rates. The mouse progressed quickly only immunohistochemistry was performed (supplementary data 1). MEMRI was performed on the second mouse and heterogeneous manganese uptake in all the visceral metastatic deposits was observed with a SE% that ranged from −0.9 to 47%. In this case the expression level of CaSR and TRPV6 receptors was markedly different, instead of what observed for the other cancer animal models. TRPV6 was highly expressed in most of the metastatic deposits with a score 4 (Figure 4) while very low-level expression of CaSR (score 0-1) was detected, except for diaphragmatic and mediastinal nodules with a CaSR score 3. All the metastatic nodules with low level of CaSR did not show any manganese uptake (Table 1) even when TRPV6 was highly expressed, while the diaphragmatic/mediastinal nodules with the higher expression level of CaSR (score 3) showed good manganese uptake with a SE% of 33-47%.

Overall, as shown in Fig. 6, the obtained results suggest a potential link between CaSR and manganese uptake. Manganese ions are analogues of calcium ions and cellular uptake of the two ions is expected to occur through the same mechanism(s) (40). However, we observed (Fig. 6) that only CaSR correlates with the cellular uptake of Mn^2+^ ions. The SE observed at MRI (i.e. 90 min after i.v. administration) could be the result of the transfer of manganese ions from the extracellular to the intracellular compartment, through the action of CaSR receptors. The absence of CaSR or its low-level expression resulted in undetectable MEMRI effects. In the liver, TRPV6 was highly expressed in both the healthy tissue and the metastatic deposits, even though normal liver tissue displayed manganese uptake while liver metastatic deposits did not.

**Figure 6.**
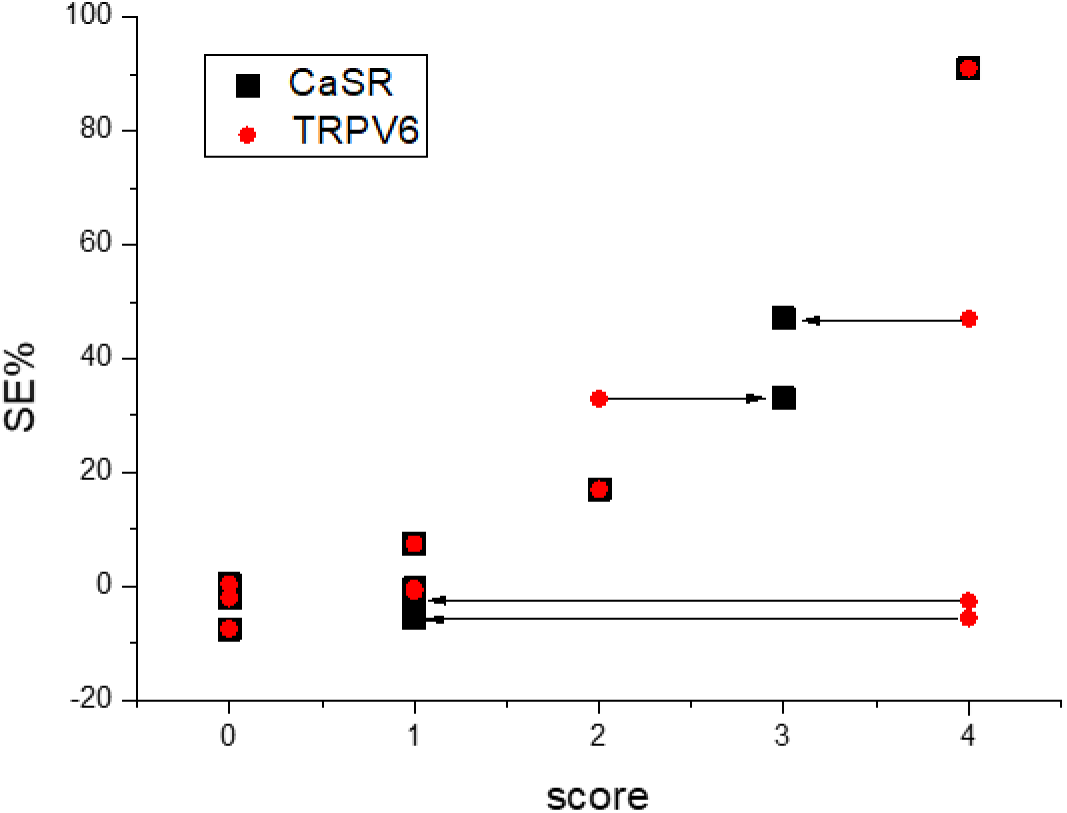
Dependence of MR signal enhancement (SE%) from the score of CaSR (black square) and TRPV6 (red circles) expression. The Mn-induced enhancement increases with CaSR expression; the expression level of TRPV6 appears not relevant for the increase of the MR signal.

One cannot exclude that other mechanisms rather than calcium receptors, are leading or affecting tumour manganese uptake. The data available from the literature, are all different and sometimes with contradictory results. Other groups explored routes to target tumour with manganese; however, the applied methods differ drastically in term of the timing between Mn^2+^ administration and MRI detection. In some studies, it was 24 hours (41), while in our experiments we observed manganese uptake at shorter times (e.g. 30, 60, 90 min until 3 hours). Another important technical aspect affecting not only tumour uptake, but also increasing the toxicity of the contrast agent, is the concentration of Mn^2+^ administered. In all experiments, we used the same concentration of manganese below the toxicity threshold and, still allowed to achieve good signal enhancement by MR (42).

Immunohistochemistry analysis carried out on the pseudometastatic human prostate cancer animal model which progressed quickly (supplementary data Figure 1), demonstrated high levels of expression of CaSR (score 4-5) either in the lung and intra-abdominal metastatic deposits. It has recently been investigated the role of CaSR in lethal prostate cancers, where higher CaSR tumour expression was associated with an approximately 2-fold higher risk for lethal progression, independently of Gleason grade and pathological stage (43). However, in prostate or breast cancer, TRPV6 is definitively associated with highly proliferative and metastatic tumours (26, 27) thus, the combination of both highly expressed calcium receptors may have affected the different timing of tumour development and progression that we observed.

Finally, we also explored the possible relationship between the manganese signal enhancement observed at MRI and other cancer biomarkers typical of these tumour cell lines, such as hormone receptor status and ki67 (supplementary data Table 2). MDA-MB-231 and PC3 cell lines demonstrated no correlation between CaSR or TRPV6 and hormone receptor or ki67 status and manganese uptake. Others applied MEMRI to investigate breast tumours and demonstrated that manganese uptake was correlated with cancer cell proliferation (41). However, in this study calcium receptors expression level was not investigated.

The mechanism behind CaSR expression and activation is the result of a very complex combination of multi-receptor actions that affect the entire cellular calcium metabolism and thus, also cancer cell proliferation. The level of extracellular calcium concentration is responsible of the activation of different intra and intercellular pathways which may affect the tumour microenvironment and, thus, the manganese uptake.

A potential limitation of this study is a control group to explore the “active role” of calcium receptors on manganese uptake by using in vivo calcium blockers, as we have performed in our previous set of experiments (15). Unfortunately, calcium blockers have important safety issues in preclinical experiments, particularly in pseudometastatic cancer animal models where their systemic administration could lead to death during the assay and the application is not allowed by the OPBA (Institutional Animal Welfare Body).

Further useful investigations aiming to demonstrate the “active role” of CaSR in MEMRI should take into consideration the knockdown/knock-in of CaSR in breast or prostate cancer cells. However, this type of experiment might very challenging due to unstable cancer features in the development of a metastatic breast or prostate cancer animal models, making the results obtained not reproducible.

To conclude the link between tumour and MEMRI is certainly interesting with high potential for future clinical application. The link tumour/calcium receptors/manganese could provide functional information related to the high potential metastatic risk of breast or prostate tumours CaSR expressing. By using quantitative MEMRI, this work has demonstrated the potential of manganese for different types of breast or prostate tumour animal models in presence of different CaSR or TRPV6 level expression. For this reasons it needs further investigation considering the different and interesting routes that manganese can use to target tumours and one of these might be the presence of calcium receptors.

## Conclusions

Manganese enhanced Magnetic Resonance Imaging (MEMRI) was able to image human breast or prostate cancer animal models with different intra- and inter-tumour expression levels of CaSR. The expression level of TRPV6 did not relate to the Mn-uptake in MEMRI experiments.

## Materials and Methods

### Human breast and prostate cancer cell lines

The human cancer cell lines PC3 (prostate cancer; ICLC) and MDA-MB-231 (breast cancer; Xenogen Corporation) were grown in RPMI 1640, with L-glutamine, 10% FCS and antibiotics (Lonza). Immunochemistry detection of CaSR and TRPV6 was performed on sections of formaldehyde-fixed paraffin-embedded cell pellets, as described below for tissue samples (data not shown).

### Human cancer animal models

In vivo experiments reviewed and approved by the OPBA (Institutional Animal Welfare Body) and by the Italian Ministry of Health with the code 237, in accordance to the National Regulation on Animal Research Resources (D.L. 116/92). Six-week old male and female NOD/SCID mice were obtained from a colony bred in house under sterile conditions. Different types of breast or prostate cancer animal models were developed, as described below, as pilot experiments to test the suitability of the models.

#### Human breast cancer orthotopic animal model

For the orthotopic breast cancer animal model, four six-week-old female, NOD-SCID mice were anesthetized with intraperitoneal injection of xylazine (10 mg/kg bw) - ketamine (100 mg/kg bw). MDA-MB-231 cells (1×10^6^ in 50 µl of serum free culture medium) were then injected in the lower left mammary fat-pad. The implantation was made under surgical sterile conditions. From 6 to 8 weeks later, when the tumour mass diameter was approximately 5 mm, as assessed by palpation and measurement with a calliper, mice were subjected to MR examination.

#### Human prostate cancer orthotopic animal model

Two six-week-old male NOD-SCID mice were anesthetized as above and PC3 cells were implanted under surgical sterile conditions. The abdomen was cleaned with iodine solution and a 1 cm midline incision was made to expose the prostate gland. One million PC3 cells suspended in 50 µl of serum free culture medium were injected into a dorsal prostatic lobe using a 30-gauge needle. After 6 weeks, the mice were subjected to MR examination and when the tumour mass diameter was approximately 5 mm, MR with manganese injection was performed.

#### Bone xenotransplant animal model of breast cancer

The osseous cancer model was set up according to Corey et al. (44). Briefly, a small hole was drilled with a 30-gauge sterile needle through the tibia plateau, with the knee flexed, of the anesthetized mouse. Using a new sterile needle fitted to a 0.3 ml sterile syringe, a single-cell suspension of 1×10^5^ MDA-MB-231 cells in 10 µl was then carefully injected in the bone marrow cavity. After 6 weeks, the mice were subjected to MR examination and when the tumour mass diameter was approximately 5 mm, MR with manganese injection was performed. The progression of osteolytic lesions was monitored by X-Ray. During radiography examination under anaesthesia, the animals were laid in prone position and left sided position on a DIGITAL Mammography (IMS, Giotto Image). Post-processing analyses were done by a dedicated workstation (Barco Mio 5MP Led).

#### Human prostate cancer animal model of metastatic disease

Prostate metastatic disease in animals was conducted as follows. Two five-week-old male, NOD-SCID mice were anesthetized with intraperitoneal injection of xylazine (10 mg/kg bw) - ketamine (100 mg/kg bw) and injected with 3×10^6^ PC3 prostate tumor cells by intracardiac route. After 6 weeks, the mice were subjected to MR examination to detect the presence of metastasis and then manganese injection was performed.

### Manganese enhanced Magnetic Resonance Imaging protocol

Magnetic resonance imaging was performed with a clinical 3T MR system (Signa EXCITE®HDxT, GE, Milwaukee, USA). The pre-clinical MR imaging protocol was developed for an experimental mouse-dedicated volume coil with a diameter of 55 mm (linear birdcage transmit/receive coil, Flick Engineering Solutions BV-General Electric-Baio G.). A specific MR protocol was applied to the cancer animal models, as previously described (15); see Table 1 for MR details parameters.

A 5 mM MnCl_2_ (Sigma Chemical Co., St. Louis, MO, USA) solution in 0.9% NaCl was prepared and 180 µl were injected intravenously per mouse (corresponding to a dose range between 7.4 mg/kg in a 24 g mouse and 8.5 mg/kg in a 21 g mouse). MR imaging was performed before and after manganese administration at the following time points 10, 30, 60 and 90 minutes, for the orthotopic cancer animal models (breast or prostate) and at 10, 30, 60, 90 minutes until 3 hours, for the metastatic cancer animal models (breast or prostate), in order to study the dynamics of manganese uptake per each metastatic lesion.

### MRI data and statistical analysis

MRI data were post-processed on a commercially available workstation (AW Volume Share 2, ADW 4.4, GE, Milano, Italy). The imaging sequences and parameters, which provided optimal signal noise to ratio (SNR) and signal enhancement (SE) relative to the manganese uptake were selected. Quantitative analyses were expressed as SE of each mouse. Tumour manganese quantification was performed by using a region of interest (ROI). The size of the region of interest depended on the diameter of the tumour (from a minimum of 2 mm to a maximum of 10 mm). The tumour SE was compared with SE of muscle and kidney, which are reference organs for CaSR and TRPV6 expression, respectively (15, 45). The SE values were divided by the background noise to yield the signal-to-noise ratio (SNR) according to the following formula:

SNR and SE% (calculated as [(SNR-SNR^0^)/ SNR^0^]×100) for all tumour lesions before and after manganese injection.

### Histopathological and immunohistochemistry analysis

Animal euthanasia was performed by the carbon dioxide method shortly after the MRI session. At necropsy tumours were excised, fixed in 10% buffered formalin and then paraffin-embedded for histological analysis according to standard techniques. The sections (3 μm) with the best representative areas of tumour tissue were identified by Hematoxylin-Eosin staining and immunostained using a BenchMark XT automated immunostainer (Ventana Medical Systems, Strasbourg, France). For bone histology, the limb was fixed in 10% buffered formalin and bones were fixed in 10% neutral buffered formalin, decalcified with a commercial chelating reagent for 2 hours and half, and trimmed for conventional paraffin embedding and histologic sectioning.

Calcium channel receptors status was performed according to previous reports (35, 36, 39). Briefly, the sections were deparaffinised and subjected to antigen-retrieval with high pH citrate buffer for 30 min. All the samples with a 1:200 dilution of the anti-calcium sensing receptor polyclonal rabbit antibody (code PA1-934A, AffinityBioReagents, Golden, CO, USA) or with buffer only, as negative control, were challenged for 30 min at 37°C. As positive staining control, parathyroid cancer was used. The antibody complex was revealed with the Polymeric System UltraView DAB Detection kit (Ventana Medical Systems). The sections were then counterstained with modified Gill’s hematoxylin, mounted in Eukitt (Bio-Optica, Milano, Italy) and observed with a light microscope (Olympus, Tokyo, Japan) using 10X, 20X, 40X and 63X objectives. A qualitative analysis of the results was performed according to intensity and pattern of the staining by two experienced pathologists (S.S. and M.T.), blind to the in vivo imaging results. A 6-point scale was used to score the intensity of the CaSR staining, going from score 0 (absent expression) to score 5 (intense, widespread expression), as described in the literature (36).

The policlonal antibody antiTRPV6 was commercially available (abcam) with cytoplasmatic localisation. Different diluitions were tested (1:500,1:1000,1: 3000) and the best one was 1:1000. The sections were deparaffinised, rehydrated and treated using the automatic immunostaininer Benchmark XT (Ventana Medical Systems, SA Strasbourg, France). Antigen retrieval was performed with citrate buffer high ph for 30 min. Then, the antibody was incubated for 1 h at 37 °C followed by addition of the polymeric detection system (Ventana Medical System Ultraview Universal DAB Detection Kit). Finally, the sections were counter-stained (automatically) with Gill’s modified haematoxylin and then cover-s lipped in Eukit. As positive tissue control, placenta was used.

## Awknowledgements

This study was supported by internal research funding assigned to the Diagnostic Imaging and Senology Unit, IRCCS Ospedale Policlinico San Martino, 16132 Genoa, Italy

## Competing interests policy

The author(s) declare no competing interests.

## In vivo MEMRI, calcium receptors expression and ki67 status

We also explored the possible relationship between the observed manganese-induced SE and other cancer biomarkers typical of these tumour cell lines, such as hormone receptor status and ki67. Immunohistochemically evaluation of Ki67, progesterone (Pgr) and estrogen (Er) receptors were performed by staining formalin fixed, paraffin-embedded 3-μm-thick tissue sections representative of the tumour previously identified through haematoxylin-eosin stained sections. The sections were deparaffinised, rehydrated and treated using the automatic immunostaininer Benchmark XT (Ventana Medical Systems, SA Strasbourg, France). Antigen retrieval was performed with citrate buffer high ph for 30 min. Then, the antibody was incubated for 1 h at 37 °C followed by addition of the polymeric detection system (Ventana Medical System Ultraview Universal DAB Detection Kit). Finally, the sections were counter-stained (automatically) with Gill’s modified haematoxylin and then cover-s lipped in Eukit. MDA-MB-231 and PC3 cell lines, are respectively, triple negative (ER, PR and her2neu) and androgen receptor negative and our results showed, no significant correlation between CaSR or TRPV6 and hormone receptor or ki67 status (see results in Table 2 supplementary data) and manganese uptake.

## Author contributions

***Conception and design:*** G.B, M.F, C.E.N. ***Development of methodology:*** G.B, M.F, C.E.N., M.C., S.S., S.A., E.G. ***Acquisition of data:*** G.B, M.F, C.E.N., F.V., F.R., L.B., S.B., S.S., L.E., M.C. ***Analysis and interpretation of data:*** G.B, M.F, C.E.N., F.R., S.S., S.A., E.G. ***Writing, review, and/or revision of the manuscript:*** G.B, M.F, C.E.N., F.R., S.S., M.C., E.G., S.A. ***Study supervision:*** G.B, M.F, C.E.N., S.A., E.G.

**Supplemental Figure 1.**
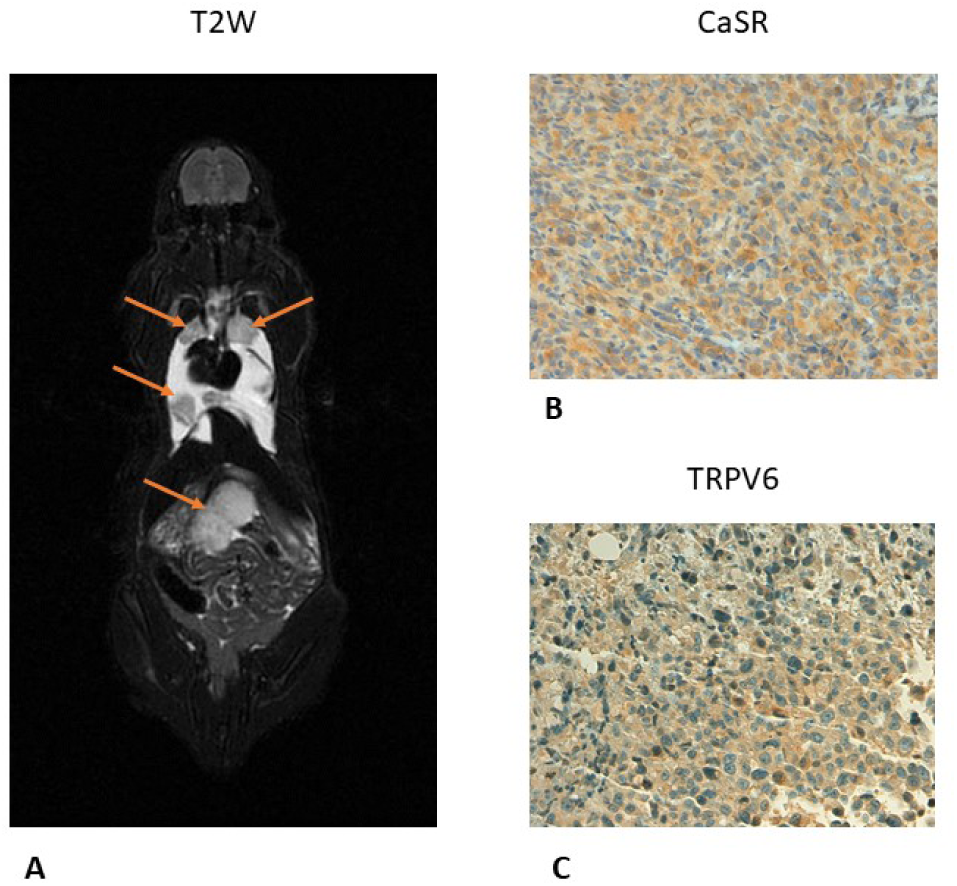
Ex-vivo MR imaging of pseudo-metastatic prostate cancer animal model and CaSR/TRPV6 receptors levels. Tumor diameter was ranging from 4 mm to 10 mm. **A.** T2-weighted image of multiple metastatic deposits within the lungs and intra-abdominal (red arrows). Bilateral pleural effusion. **BC.** Immunohistochemistry (IHC) of CaSR and TRPV6 receptors displayed intense staining (score 4) in all the metastatic deposits in keeping with high aggressive prostate cancer.

**Supplemental Figure 2.**
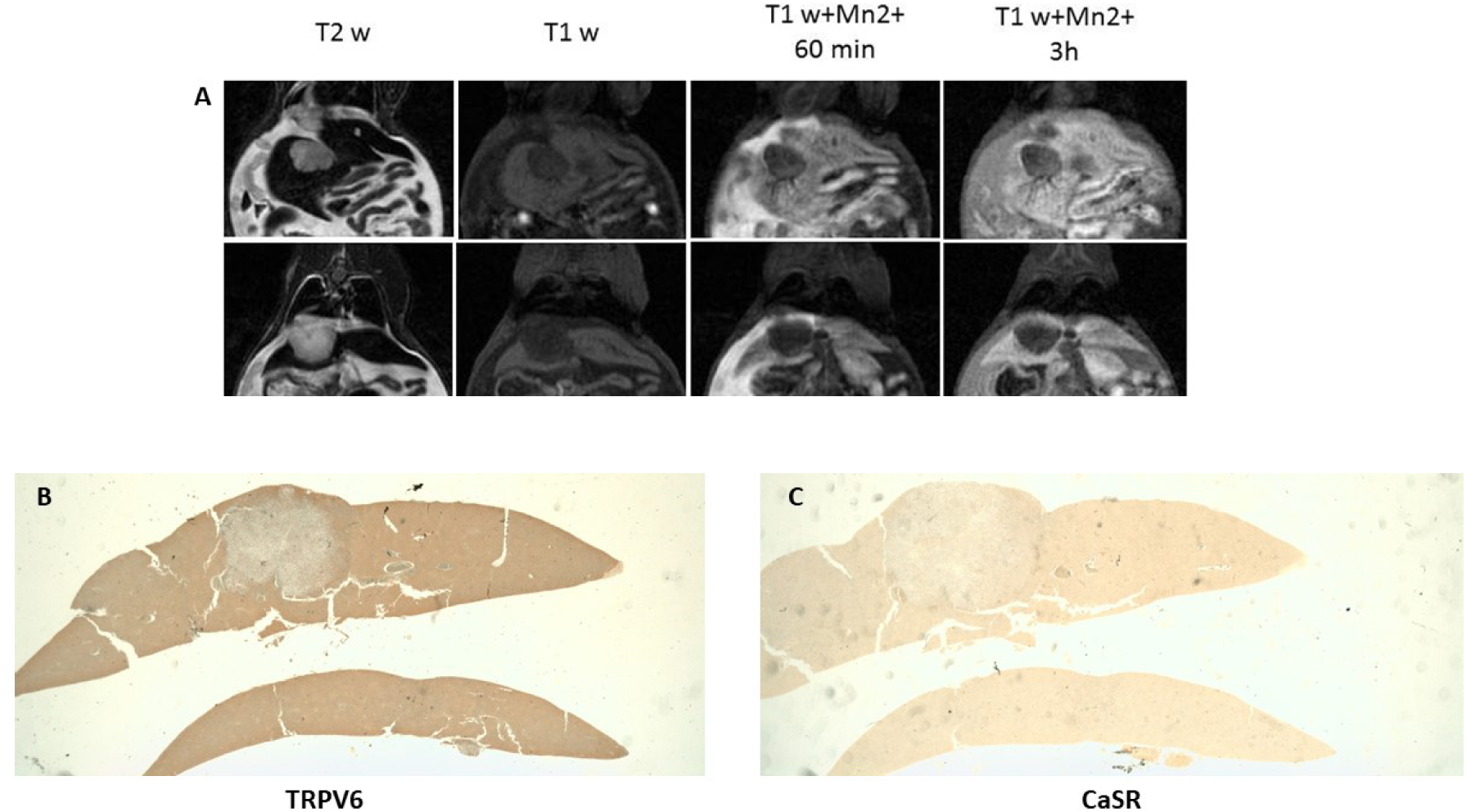
Manganese enhanced MR imaging (MEMRI) of pseudo-metastatic prostate cancer animal model and CaSR/TRPV6 receptors levels. **A.** T2-weighted image of a liver metastasis. T1-weighted gradient echo images (T1 WI) recorded before and after 60 minutes and 3 hours of Mn2+ administration. No significant manganese uptake is appreciated, at both 60 minutes or 3 hours after Mn2+ administration. **BC.** Immunohistochemistry (IHC) of TRPV6 and CaSR: TRPV6 receptors displayed intense staining (score 4), while a rare positive staining of CaSR was detected in tumour cells (score 1). TRPV6 expression level in normal liver tissue was higher compared to CaSR expression level.

## References

1. Berridge, M.J, Bootman, M.D, Roderick, H.L. Calcium signalling: dynamics, homeostasis and remodelling, Nat Rev Mol. Cell Biol. 2003; 4: 517–529.

2. Berridge, M.J, Lipp, P. Bootman, M.D. The versatility and universality of calcium signalling, Nat. Rev. Mol. Cell Biol. 2000; 1: 11–21.

3. Clapham, D.E. Calcium signaling, Cell 2007; 131: 1047–1058.

4. Machaca, K. Ca (2+) signaling, genes and the cell cycle, Cell Calcium 2011; 49: 323–330.

5. Parkash, J. Asotra, K. Calcium wave signaling in cancer cells, Life Sci. 2010; 87: 587–595.

6. Missiaen L, Robberecht W, van den Bosch L, Callewaert G, Parys JB, Wuytack F., et al Abnormal intracellular Ca (2+) homeostasis and disease, Cell Calcium 2000; 28: 1–21.

7. Roderick, H.L. Cook, S.J. Ca2+ signalling checkpoints in cancer: remodelling Ca2+ for cancer cell proliferation and survival, Nat. Rev. Cancer 2008; 8: 361–375.

8. Wen, L. Shi, X. He, L. Han, D. Manganese-Enhanced Magnetic Resonance Imaging for Detection and Characterization of Colorectal Cancers. Tomography 2018; 4: 78–83.

9. Bianchi A, Gobbo OL, Dufort S, Sancey L, Lux F, Tillement O et al. Orotracheal manganese-enhanced MRI (MEMRI): An effective approach for lung tumor detection. NMR Biomed. 2017; 30: 11.

10. Gianolio E, Arena F, Di Gregorio E, Pagliarin R, Delbianco M, Baio G et al. MEMRI and tumors: a method for the evaluation of the contribution of Mn(II) ions in the extracellular compartment. NMR Biomed 2015; 28: 1104–1110.

11. Lin, Y.J. Koretsky, A.P. Manganese ion enhances T1-weighted MRI during brain activation: An approach to direct imaging of brain function. Magn Reson Med 1997; 38: 378– 388.

12. Pautler, R.G. Silva A.C. Koretsky, A.P. In vivo neuronal tract tracing using manganese-enhanced magnetic resonance imaging. Magn Reson Med 1998; 40: 740–748.

13. Pautler, R.G. Biological applications of manganese-enhanced magnetic resonance imaging. Methods Mol Med 2006; 124: 365–386.

14. Silva, A.C. Lee, J.H. Aoki, I. Koretsky, A.P. Manganese-enhanced magnetic resonance imaging (MEMRI): Methodological and practical considerations. NMR Biomed 2004; 17: 532–543.

15. Baio G, Fabbi M, Emionite L, Cilli M, Salvi S, Ghedin P, et al, In vivo imaging of human breast cancer mouse model with high level expression of calcium sensing receptor at 3T. Eur Radiol 2012; 22: 551–558.

16. Brennan SC, Thiem U, Roth S, Aggarwal A, Fetahu ISh, Tennakoon S, et al. Calcium sensing receptor signalling in physiology and cancer. Biochim Biophys Acta. 2013; 1833: 1732–1744.

17. Li X, Li L, Moran MS, Jiang L, Kong X, Zhang H. et al. Prognostic significance of calcium-sensing receptor in breast cancer. Tumour Biol. 2014; 35:5709–5715.

18. Aggarwal A, Prinz-Wohlgenannt M, Tennakoon S, Höbaus J, Boudot C, Mentaverri R et al. The calcium-sensing receptor: A promising target for prevention of colorectal cancer. Biochim Biophys Acta 2015; 1853: 2158–2167.

19. Joeckel E, Haber T, Prawitt D, Junker K, Hampel C, Thüroff JW. et al. High calcium concentration in bones promotes bone metastasis in renal cell carcinomas expressing calcium-sensing receptor. Mol Cancer 2014; 28: 13:42.

20. Shapovalov, G. Ritaine A., Skryma R. Prevarskaya N. Role of TRP ion channels in cancer and tumorigenesis. Semin Immunopathol 2016; 38: 357–369.

21. Monteith, G.R. McAndrew, D. Faddy, H.M. Roberts-Thomson, S.J. Calcium and cancer: targeting Ca2+ transport. Nat Rev Cancer 2007; 7: 519–530.

22. Roderick, H.L. Cook, S.J. Ca2+ signalling checkpoints in cancer: remodelling Ca2+ for cancer cell proliferation and survival. Nat Rev Cancer 2008; 8: 361–375.

23. Stoerger, C. Flockerzi, V. The transient receptor potential cation channel subfamily V member 6 (TRPV6): genetics, biochemical properties, and functions of exceptional calcium channel proteins. Biochem Cell Biol. 2014; 92: 441–448.

24. Fecher-Trost, C. Weissgerber, P. Wissenbach, U. TRPV6 channels. Handb Exp Pharmacol 222:359–384 (2014).

24. Bodding, M. Flockerzi, V. Ca2+ dependence of the Ca2 + −selective TRPV6 channel. J Biol Chem 2004; 279: 36546–36552.

25. Bodding, M. Wissenbach, U. Flockerzi, V. The recombinant human TRPV6 channel functions as Ca2+ sensor in human embryonic kidney and rat basophilic leukemia cells. J Biol Chem 2002; 277: 36656–36664.

26. Bolanz, K.A. Hediger, M.A. Landowski, C.P. The role of TRPV6 in breast carcinogenesis. Mol Cancer Ther 2008; 7: 271–279.

27. Zhuang L1, Peng JB, Tou L, Takanaga H, Adam RM, Hediger MA, et al. Calcium-selective ion channel, CaT1, is apically localized in gastrointestinal tract epithelia and is aberrantly expressed in human malignancies. Lab Invest 2002; 82: 1755–1764.

28. Peng JB, Zhuang L, Berger UV, Adam RM, Williams BJ, Brown EM, et al. CaT1 expression correlates with tumor grade in prostate cancer. Biochem Biophys Res Commun 2001; 282: 729–734.

29. Wissenbach U, Niemeyer BA, Fixemer T, Schneidewind A, Trost C, Cavalie A, et al Expression of CaT-like, a novel calcium-selective channel, correlates with the malignancy of prostate cancer. J Biol Chem 2001; 276: 19461–19468.

30. Bodding, M. TRP proteins and cancer. Cell Signal 2007; 19: 617–624.

31. Peters AA, Simpson PT, Bassett JJ, Lee JM, Da Silva L, Reid LE, et al. Calcium channel TRPV6 as a potential therapeutic target in estrogen receptor-negative breast cancer. Mol Cancer Ther 2012; 11: 2158–2168.

32. Corey E, Quinn JE, Bladou F, Brown LG, Roudier MP, Brown JM. et al. Establishment and characterization of osseous prostate cancer models: intra-tibial injection of human prostate cancer cells. Prostate 2002; 52: 20–33.

33. Prevarskaya, N. Skryma, R. Shuba, Y. Calcium in tumour metastasis: new roles for known actors. Nat Rev Cancer 2011; 11: 609–618.

34. Yang, S.L. Cao, Q. Zhou, K.C. Feng, Y.J. Wang, Y.Z. Transient receptor potential channel C3 contributes to the progression of human ovarian cancer. Oncogene 2009; 28: 1320–1328.

35. Prevarskaya, N., Zhang, L., Barritt G. TRP channels in cancer. Biochim Biophys Acta 2007; 1772: 937–946.

36. Mihai R, Stevens J, McKinney C, Ibrahim NB et al. Expression of the calcium receptor in human breast cancer-a potential new marker predicting the risk of bone metastases. Eur J Surg Oncol 2006; 32: 511–515.

37. Baio G, Rescinito G, Rosa F, Pace D, Boccardo S, Basso L, et al. Correlation between Choline Peak at MR Spectroscopy and Calcium-Sensing Receptor Expression Level in Breast Cancer: A Preliminary Clinical Study. Mol Imaging Biol. 2015; 17: 548–556.

37. Parkash, J. Chaudhry, MA. Rhoten, WB. Calbindin-D28k and calcium sensing receptor cooperate in MCF-7 human breast cancer cells. Int J Oncol. 2004; 24: 1111–1119.

38. Parkash, J. Chaudhry, MA. Rhoten, WB. Ca(2+) sensing receptor activation by CaCl(2) increases [Ca2+]i resulting in enhanced spatial interactions with calbindin-D28k protein. Int J Mol Med. 2004; 13: 3–11.

39. Castets CR, Koonjoo N, Hertanu A, Voisin P, Franconi JM, Miraux S et al. In vivo MEMRI characterization of brain metastases using a 3D Look-Locker T1-mapping sequence. Sci Rep 2016; 6: 39449.

40. Merritt, J. E. Jacob, R. Hallam, T.J. Use of Manganese to Discriminate between Calcium Influx and Mobilization from Internal Stores in Stimulated Human Neutrophils. J Biol Chem 1989; 264: 1522–1527.

41. Nofiele, J.T. Czarnota, G.J. Cheng, H.L. Noninvasive manganese-enhanced magnetic resonance imaging for early detection of breast cancer metastatic potential. Mol Imaging. 2014; 13. doi: 10.2310/7290.2013.00071.

42. Grünecker B, Kaltwasser SF, Peterse Y, Sämann PG, Schmidt MV, Wotjak CT, et al. Fractionated manganese injections: effects on MRI contrast enhancement and physiological measures in C57BL/6 mice. NMR Biomed 2010; 23: 913–921.

43. Ahearn TU, Tchrakian N, Wilson KM, Lis R, Nuttall E, Sesso HD et al Calcium-Sensing Receptor Tumor Expression and Lethal Prostate Cancer Progression. Clin Endocrinol Metab 2016; 101: 2520–2527.

44. Hoenderop JG, Vennekens R, Müller D, Prenen J, Droogmans G, Bindels RJ, et al. Function and expression of the epithelial Ca(2+) channel family: comparison of mammalian ECaC1 and 2. J Physiol 2001; 537: 747–761.

